# Pharmacological modulation of AMPA receptor surface diffusion restores hippocampal synaptic plasticity and memory in Huntington’s disease

**DOI:** 10.1101/297069

**Authors:** Hongyu Zhang, Chunlei Zhang, Jean Vincent, Diana Zala, Caroline Benstaali, Matthieu Sainlos, Dolors Grillo-Bosch, Yoon Cho, Denis J. David, Frederic Saudou, Yann Humeau, Daniel Choquet

**Author notes:** Correspondence should be addressed to D.C., Y.H. or H.Z.

## Abstract

Impaired hippocampal synaptic plasticity is increasingly considered to play an important role in cognitive impairment in Huntington’s disease (HD). However, the molecular basis of synaptic plasticity defects is not fully understood. Combining live-cell nanoparticle tracking and super-resolution imaging, we show that dysregulation of AMPA receptors (AMPARs) surface diffusion represents a molecular basis underlying the aberrant hippocampal synaptic plasticity during HD. AMPARs surface diffusion is increased in various HD neuronal models, which results in the failure of AMPARs surface stabilization after long-term potentiation (LTP) stimuli. This appears to result from a defective brain-derived neurotrophic factor (BDNF) - tyrosine receptor kinase B (TrkB) - Ca2+/calmodulin-dependent protein kinase II (CaMKII) signaling pathway that impacts the interaction between the AMPAR auxiliary subunit stargazin and postsynaptic density protein 95 (PSD-95). Notably, the disturbed AMPAR surface diffusion is rescued, via BDNF signaling pathway and by the antidepressant tianeptine. Tianeptine also restores the impaired LTP and hippocampus-dependent memory as well as anxiety/depression-like behavior in different HD mouse models. We thus unveil a mechanistic framework underlying hippocampal synaptic and memory dysfunction and propose a new perspective in HD treatment by targeting AMPAR surface diffusion.

---

Cognitive deficits and psychiatric disturbance prior to motor dysfunction have been widely documented in preclinical Huntington’s disease (HD) gene carriers ^1,2^. These manifestations have traditionally been attributed to degeneration or death of corticostriatal neurons ^3^. However, mounting evidence points to the involvement of deficits in hippocampal synaptic plasticity. This is supported by the findings that hippocampal long-term potentiation (LTP), a major form of synaptic plasticity widely regarded as a molecular basis for learning and memory, is greatly impaired in different categories of HD mouse models at pre- or early-symptomatic stage ^4–7^. Moreover, the abnormally regained ability to support long-term depression (LTD) has also been reported in HD mice ^8^. Consistently, behavioral studies reveal deterioration of hippocampal-associated spatial memory in distinct HD murine models ^7,9,10^, primate model ^11^, and patients ^12^.

The molecular mechanisms underlying hippocampal synaptic and memory dysfunctions are not well understood but the BDNF signaling pathway seems to play an important role. BDNF is a potent, positive modulator of LTP ^13^. The down-regulation of its protein production ^6,14^ and the imbalance between the expression of its high-affinity TrkB receptor ^15,16^ and low-affinity p75 neurotrophin receptor (P75^NTR^) ^9,17-19^ have been implicated in the hippocampal synaptic and memory defects in HD. Indeed, administration of BDNF or P75^NTR^ gene knockdown ameliorates HD-associated synaptic and memory dysfunction ^6,9^. However, the signaling mechanisms mediating BDNF modulation of synaptic plasticity and mechanism-based pharmacological treatment strategies remain largely unexplored. This may have significant therapeutic implications as the application of exogenous BDNF is not clinically practical due to its instability in the bloodstream and its inability to cross the blood-brain barrier ^20,21^, and genetic intervention on human subjects may carry ethical issues.

AMPA receptors (AMPARs) are the major excitatory neurotransmitter receptors. The regulated trafficking of AMPARs to and from the synapses is thought to be a key mechanism underlying glutamatergic synaptic plasticity ^22–24^. Animal studies reveal that AMPAR trafficking plays a pivotal role in experience-driven synaptic plasticity and modification of behavior ^25^. In pathological conditions, acute stress or response to stress hormones (eg, noradrenaline, corticosterone) alters AMPAR trafficking and memory encoding processes ^26–28^. Thus, monitoring and manipulating synaptic AMPAR trafficking emerges as a useful tool to study cognitive function and dysfunction in animal models. Synaptic delivery of AMPAR involves intracellular trafficking, insertion to the plasma membrane by exocytosis, and lateral diffusion at the neuronal surface ^23,29^. For many years, endo/exocytosis have been considered to be the main routes for exit and entry of receptors from and to postsynaptic sites, respectively. However, our lab and others have established in the last decade that receptor surface diffusion is a key step for modifying receptor numbers at synapses ^22,30,31^. We have demonstrated that deregulated AMPAR surface diffusion primarily contributes to the impaired LTP in stress/depression models^28^. Most importantly, we found that AMPAR surface diffusion can be pharmacologically modulated by a clinically used antidepressant tianeptine (S 1574, [3-chloro-6-methyl-5, 5-dioxo-6,11-dihydro-(c,f)-dibenzo-(1,2-thiazepine)-11-yl) amino]-7 heptanoic acid), which restores impaired LTP in the stress/depression model ^32^. As impaired synaptic plasticity is a common mechanism underlying both cognitive impairment and psychiatric disturbance such as anxiety and depression ^6,9,24,27,33^, the major early-onset symptoms in HD ^1,2,34^, here we have examined whether AMPAR surface diffusion is disturbed in HD models, how this is linked with impaired BDNF signaling and whether pharmacological modulation of AMPAR surface diffusion by tianeptine can serve as a promising therapeutic strategy to improve synaptic and memory dysfunction as well as anxiety/depression behavior in HD.

## RESULTS

### Increased AMPAR surface diffusion in three different HD cellular models

AMPARs are heteromeric proteins composed of different combinations of GluA1, GluA2, GluA3 or GluA4 subunits, in which GluA1-GluA2 di-heteromers are the most common combination in adult neurons. We thus first investigated endogenous GluA2-AMPAR surface diffusion using the single nanoparticle tracking approach in which a Quantum Dot (QD) is coupled to an antibody specific for the extracellular domain of the endogenous GluA2 subunit (Fig. 1a) ^32^. We initially used rat primary hippocampal neuronal cultures transfected with exon1 mutant huntingtin which contains 69 polyglutamine expansion (exon1-polyQ-HTT), with exon1 wild-type huntingtin with 17 polyglutamine (exon1-wHTT) and empty vector as controls ^35^. Compared to empty vector and exon1-wHTT, expression of exon1-polyQ-HTT significantly increased the surface diffusion of GluA2-AMPAR (Fig. 1b, c top panel). To avoid possible transfection artifacts, we next used primary hippocampal neurons from male R6/1 heterozygous transgenic mice, which overexpress the first exon of human HTT with 115 polyQ and represent a fast model of HD. Only male HD mice were used throughout the paper in order to eliminate the possible influence of gender differences ^34^. Similarly, an increase in GluA2-AMPARs surface diffusion was observed in neurons from R6/1 mice compared to WT littermate controls (Fig.1d top panel). Furthermore, to circumvent overexpression artifacts and to better mimic the genetic situation in patients, we used neurons from male homozygous *Hdh*^Q111/Q111^ knock-in mouse, in which polyQ repeats are directly engineered into the mouse HTT genomic locus and wHTT/polyQ-HTT is expressed at endogenous levels. Consistently, neurons from *Hdh*^Q111/Q111^ knock-in mouse also displayed marked increase in GluA2 surface diffusion compared to WT littermates (Fig. 1e, top panel). These changes were partially due to a decreased fraction of immobile GluA2-AMPAR (surface diffusion ≤ 0.01 µm^2^/s) (Fig. 1c-e, bottom panels). Cumulative distributions of diffusion coefficients shift towards the right in the 3 HD models, indicating an increased GluA2-AMPAR surface diffusion.

**Figure 1.**
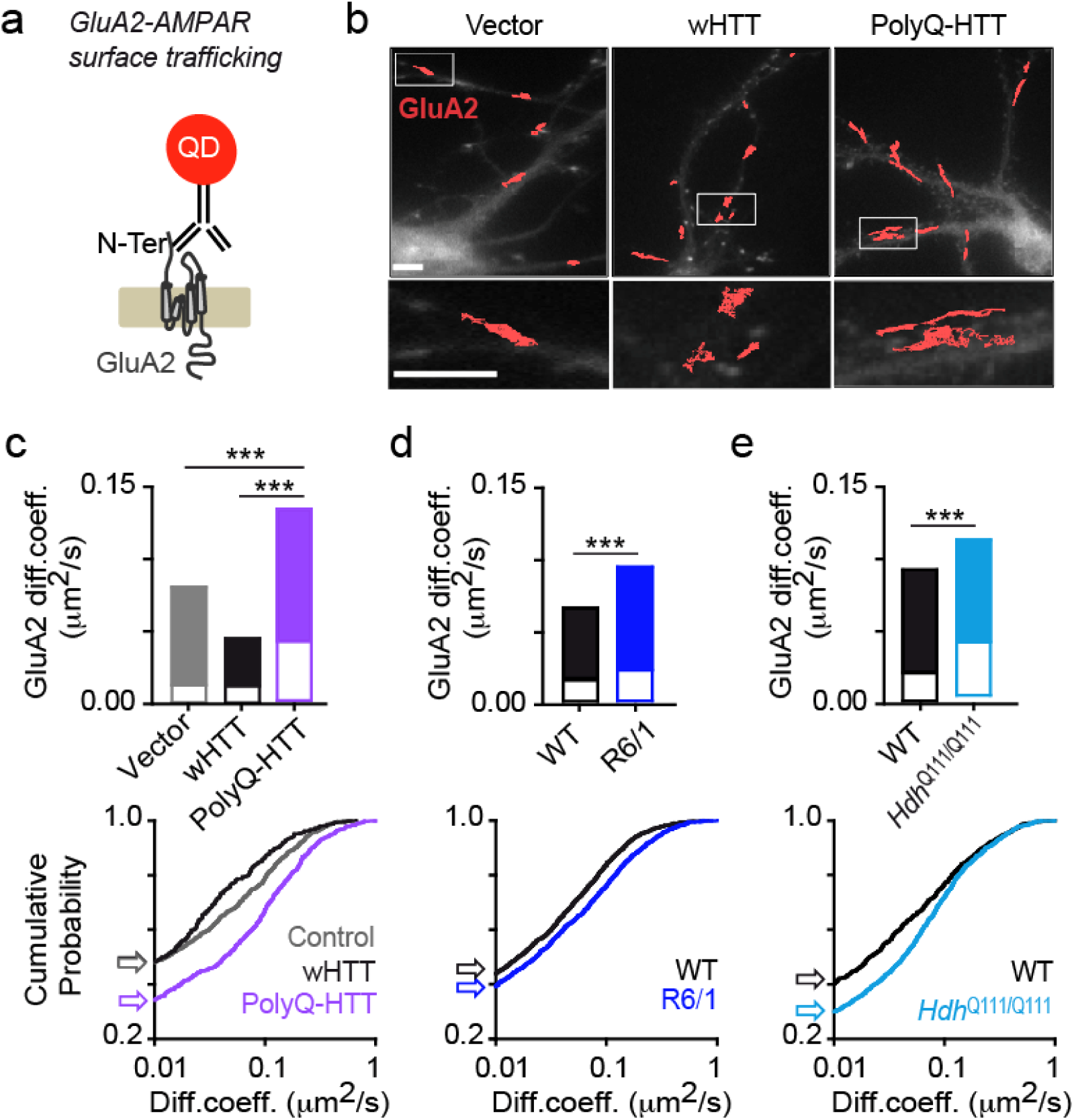
Deregulated GluA2-AMPAR surface diffusion in different complementary HD cellular models **(a)** Experimental scheme showing that for endogenous GluA2-AMPAR surface tracking, hippocampal neurons were incubated with mouse monoclonal antibody against N-terminal extracellular domain of GluA2 subunit followed by QD anti-mouse IgG. (**b**) Typical GluA2-QD trajectories (red) in hippocampal neurons expressing vector, exon1-wHTT and exon1-polyQ-HTT, respectively. Lower panels represent enlarged GluA2-QD trajectories. Scale bars, 10μm. **(c, d, e)** Top panels, GluA2-AMPAR diffusion coefficients (median ± 25-75% interquartile range (IQR)) in rat hippocampal neurons expressing vector, exon1-wHTT, and exon1-polyQ-HTT; n = 844, 382 and 695 trajectories, respectively **(c)**, in hippocampal neurons from R6/1 mice and WT littermate controls; n =1885 and 1994 trajectories, respectively **(d),** and in hippocampal neurons from *Hdh*^Q111/Q111^ mice and WT littermate controls; n =1571 and 886 trajectories, respectively **(e)**. Bottom panels, cumulative probability of GluA2 diffusion coefficient of respective top panel. The first point of the probability corresponding to the fraction of immobile receptors with diffusion coefficients ≤ 0.01 μm2/s was showed by arrows. Note that the cumulative curve shifts toward right indicating an increased GluA2 surface diffusion. Significance was determined by Kruskal-Wallis test followed by Dunn’s Multiple Comparison Test (**c**), or Mann-Whitney test (**d, e**). \*\*\**P* < 0.001.

As AMPARs are composed of different subunits, which can be differentially trafficked, we also investigated the surface diffusion of the GluA1-AMPAR population in the 4^th^ cellular model of HD, in which full-length (FL) HTT with 75 polyQ and FL-wHTT with 17Q were overexpressed in rat hippocampal neurons. Similarly, the surface diffusion coefficients of GluA1-AMPARs were markedly increased in neurons expressing FL-polyQ-HTT compared to neurons expressing FL-wHTT (Supplemental Fig. 1). Collectively, these data demonstrate in different complementary cellular models of HD that the surface diffusion of GluA2 and GluA1 subunits of AMPAR was markedly increased.

### Surface GluA2-AMPARs failed to stabilize on the neuronal surface after chemical LTP (cLTP) induction in an HD cellular model

The increased AMPAR surface diffusion in basal conditions prompted us to ask whether this could potentially lead to abnormal AMPAR surface stabilization during activity-dependent synaptic plasticity, such as LTP. Indeed, on the one hand, it has been shown that activity-dependent synaptic potentiation is associated with immobilization and subsequent accumulation of AMPARs at synapses ^31,36,37^. On the other hand, polyQ expansion of HTT is associated with impaired LTP ^4–7^. We thus examined AMPAR surface diffusion before and after three-minute cLTP stimuli ^38^ (300μM Glycine, 1μM Picrotoxin, without Mg^2+^) in rat hippocampal neurons overexpressing FL-wHTT or FL-polyQ-HTT using a super-resolution imaging method, Universal Point Accumulation Imaging in Nanoscale Topography (uPAINT) ^39^. uPAINT is not only able to generate super-resolved images but also provides dynamic information with large statistics revealing localization-specific diffusion properties of membrane biomolecules. Endogenous GluA2-AMPARs were tracked with ATTO 647 labeled anti-extracellular GluA2 antibody and sorted into two groups according to their diffusion coefficient (immobile, Log (D) =< -2; mobile, Log (D) > -2). In FL-wHTT-expressing neurons, we observed a decrease in the ratio of mobile to immobile AMPAR after cLTP stimuli relative to basal condition (Pre-LTP), reflecting an immobilization of surface AMPARs (Fig.2a). In contrast, the ratio of mobile to immobile AMPAR in FL-polyQ-HTT-expressing neurons was not significantly different before and after LTP stimuli (Fig.2b). These data suggest that AMPARs fail to stabilize at the neuronal surface after LTP stimuli in HD models. This could explain, at least in part, the defects in the potentiation of AMPAR-mediated synaptic transmission in HD.

**Figure 2.**
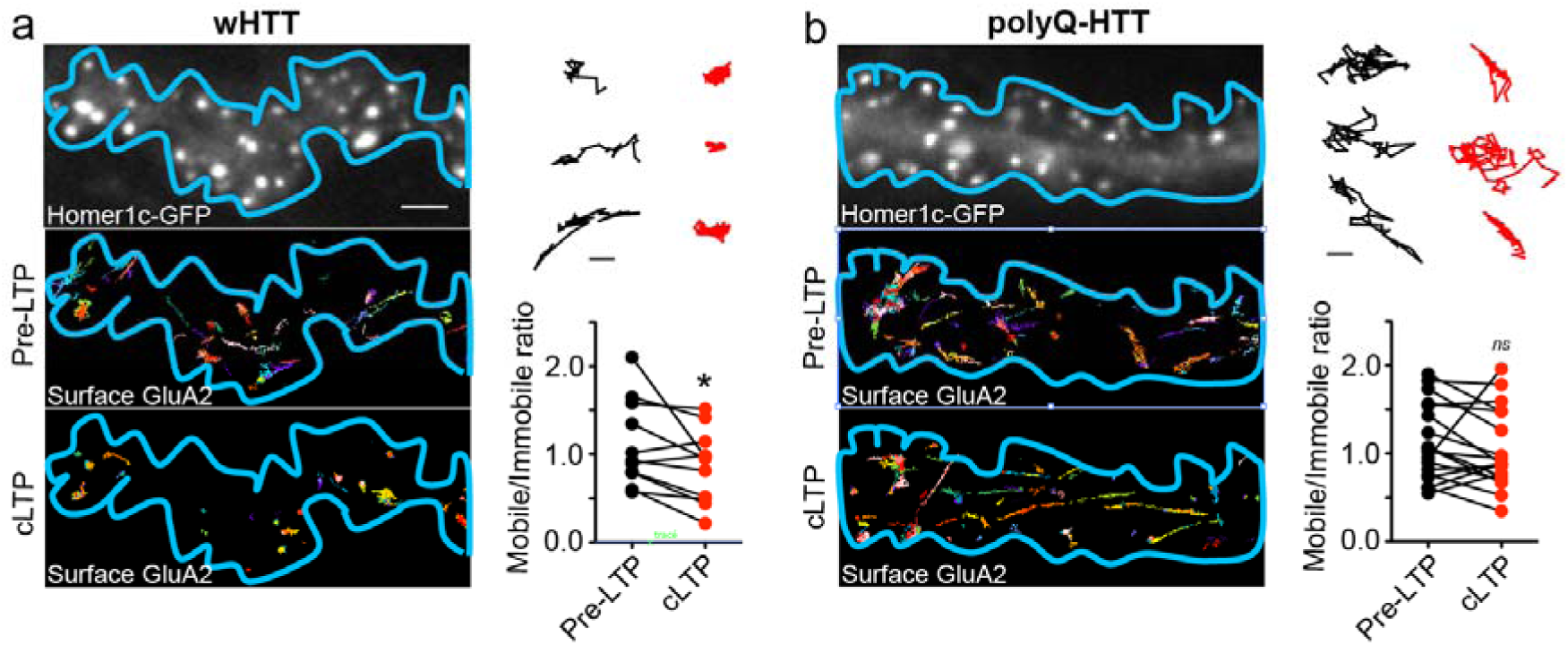
GluA2-AMPAR in polyQ-HTT-expressing neurons failed to stabilize on the neuronal surface after chemical LTP (cLTP) stimulation. (**a, b**) Top-left panels, epifluorescence image of a dendritic segment co-expressing Homer1c-EGFP (synaptic marker) and FL-wHTT (**a**) or FL-polyQ-HTT **(b**); middle- and bottom-left panels, corresponding super-resolution image of endogenous GluA2-AMPAR trajectories accumulated from 2000 images before (middle-left panels) and after cLTP stimulation (bottom-left panels) for the outlined region in the epifluorescence image. Scale bars, 10μm. Top-right panels, enlarged typical GluA2-AMPAR trajectories before (black) and after cLTP induction (red) in FL-wHTT-(**a**) and FL-polyQ-HTT-expressing neurons (**b**). Scale bars, 10μm. Bottom-right panels, the ratio of mobile to immobile fraction of the diffusion coefficient (D) before and after cLTP induction in FL-wHTT-(**a**) and FL-polyQ-HTT-expressing neurons (**b**). Immobile fraction was identified as the proportion of receptors with D ≤ 0.01 μm2/s while mobile fraction with D > 0.01 μm2/s. Paired t-test was used. \**P* < 0.05; *ns*, not significant.

### Impaired BDNF-TrkB-CaMKII signaling through modulation of the interaction between stargazin and PSD95 contributes to the deregulation of AMPAR surface diffusion in the hippocampus

BDNF is a prominent positive modulator of LTP ^13^, which has been proposed to induce the delivery of AMPARs to the synapse under basal conditions ^40^. However, it is not known whether and how AMPAR surface diffusion is modulated by BDNF signaling and whether it plays a role in HD pathogenesis. We thus asked if deficient BDNF signaling could account for the aberrant AMPAR surface diffusion in HD mouse models. We first characterized changes in the protein level of BDNF in HD mice. Consistent with a previous report ^6^, using ELISA, we observed a significant decrease in the protein level of BDNF in the hippocampus of 10-week-old R6/1 and *Hdh*^Q111/Q111^ mice compared to respective littermate controls (Fig. 3b). We next studied BDNF intracellular transport in 3 complementary HD cellular models, as data on the BDNF intracellular transport in the hippocampus of HD mice are still lacking. Slower anterograde and retrograde BDNF intracellular transport was exhibited by neurons expressing polyQ-HTT relative to wHTT-expressing neurons (Fig.3c, 3d), and by R6/1 and *Hdh*^Q111/Q111^ mouse hippocampal neurons compared to respective WT littermate controls (Fig. 3e and 3f, respectively). Note that we observed slower BDNF velocity in neurites (Fig. 3d, 3e) than in axons in hippocampal neurons (Fig. 3f), which is consistent with previous studies in cortical neurons ^41,42^. Altogether, these data suggested that reduced BDNF protein production and impaired intracellular transport are common features to different categories of HD models.

Next we dissected the potential signaling mechanism by which BDNF modulates AMPAR surface diffusion in HD models. BDNF is known to bind to TrkB receptors, leading to the activation of CaMKII^13^, which is critically required for the synaptic recruitment of AMPAR during both development and plasticity ^36^. Active CaMKII phosphorylated at threonine 286 (T286) is reported to be reduced in the hippocampus of *Hdh*^Q111/Q111^ mouse models ^9^. We next confirmed the decrease in CaMKII activity in a HD cellular model by co-transfecting rat hippocampal neurons with FL-wHTT/FL-polyQHTT and a fluorescence resonance energy transfer (FRET)-based CaMKII**α**, named REACH-CaMKII. The amino and carboxy termini of REACH-CaMKII are labeled with the FRET pair of monomeric enhanced green fluorescent protein (mEGFP) and resonance energy-accepting chromoprotein (REACh), a non-radiative yellow fluorescent protein variant ^43^. The activation of REACh-CamKII associated with T286 phosphorylation changes the conformation of CaMKII**α** to the open state in which its kinase domain is exposed, thereby decreasing FRET and increasing the fluorescence lifetime of mEGFP (Supplemental Fig. 2a). Rat hippocampal neurons transfected with GFP-PSD95 alone were used as negative control. GFP-PSD95-expressing cells showed high lifetime in both dendritic puncta and shaft indicating no FRET. FL-wHTT- and FL-polyQ-HTT-expressing cells both exhibited FRET revealed by shorter lifetime than GFP-PSD95-expressing cells, indicating activated CaMKIIα. However, the REACh-CaMKII**α** lifetime in dendritic puncta (Supplemental Fig. 2c) and shaft (Supplemental Fig. 2d) in FL-polyQ-HTT-expressing cells are significantly lower than in FL-wHTT-expressing cells, indicating stronger FRET and thus weaker CaMKII**α** activity.

**Figure 3.**
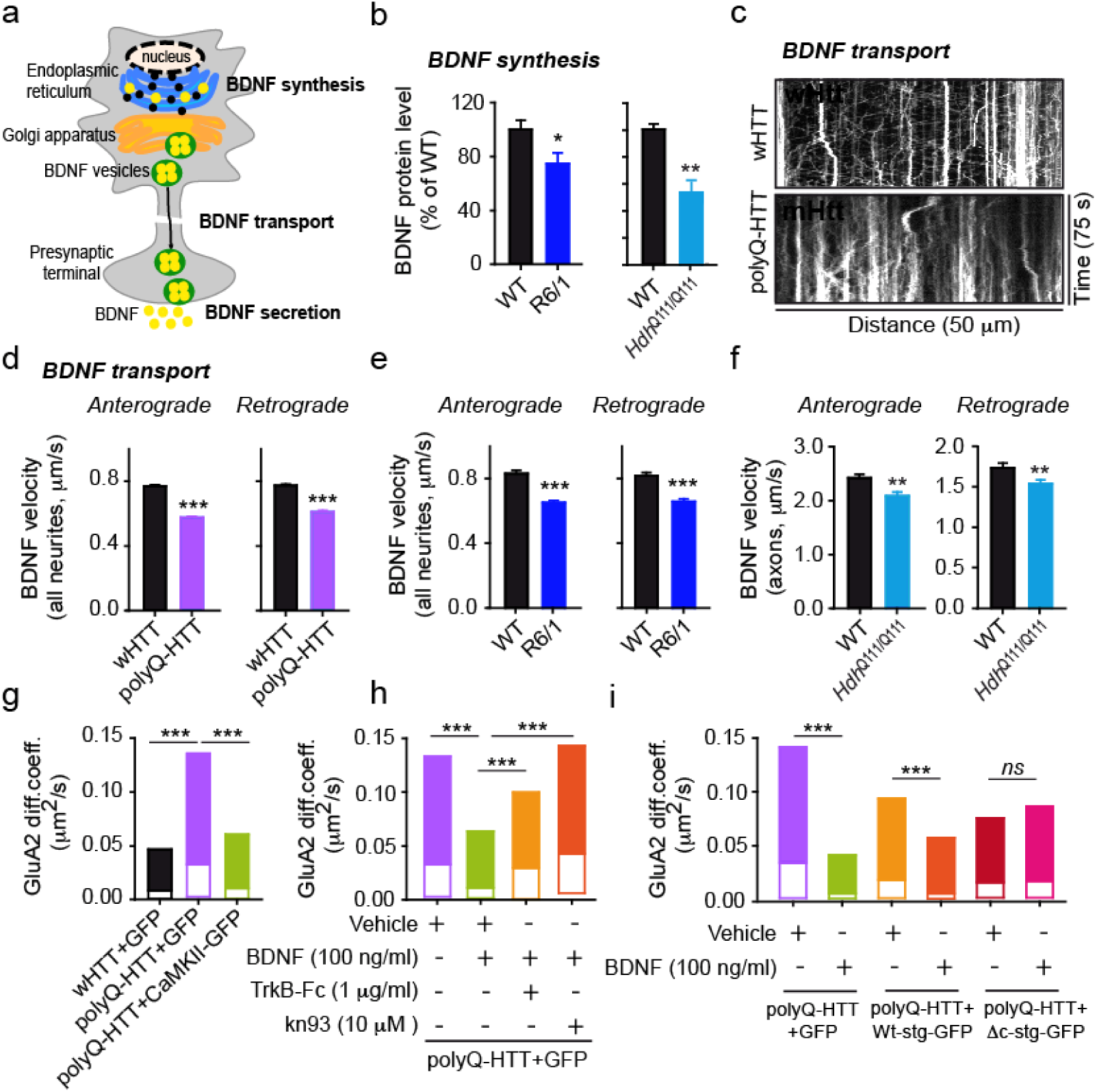
Impaired BDNF-TrkB-CaMKII signaling through the interaction between stargazin and PSD95 contributes to the deregulation of AMPAR surface diffusion in HD models (**a**) Schematic diagram showing that BDNF can be modulated at synthesis, transport and secretion level. (**b**) Hippocampal BDNF protein level determined by ELISA in R6/1 and *Hdh*^Q111/Q111^ mice; values are mean ± s.e.m (% of WT); n = 21 and 14 mice for WT and R6/1; n = 6 and 9 mice for WT and *Hdh*^Q111/Q111^, respectively. (**c**) Representative kymographs of intracellular transport of BDNF-containing vesicles (white trajectories) in a neurite (50 μm from soma) over 75 seconds (s) in wHTT- and polyQ-HTT-expressing rat hippocampal neurons. The velocity of BDNF transport was reflected by the slope of trajectories (moving distance against time). (**d, e, f**) Anterograde and retrograde BDNF transport velocity in all neurites of wHTT- and polyQ-HTT-expressing rat hippocampal neurons (**d**), and hippocampal neurons from R6/1 mouse line (**e**), and in the axon of hippocampal neurons from *Hdh*^Q111/Q111^ mouse line (**f**); values are mean ± s.e.m; n = 5569, 5656, 5227 and 5706 trajectories for anterograde and retrograde wHTT and polyQ-HTT, respectively; n = 1424, 1710, 1376, and 1487 trajectories for anterograde and retrograde WT and R6/1, respectively; n = 236, 261, 194 and 256 trajectories for anterograde and retrograde WT and *Hdh*^Q111/Q111^, respectively. (**g, h, i**) GluA2-AMPAR diffusion coefficients in rat hippocampal neurons co-expressing FL-wHTT/polyQ-HTT and GFP, or FL-polyQ-HTT and CamKII-GFP; n = 656, 685, and 349 trajectories, respectively (**g**), in neurons co-expressing FL-polyQ-HTT and GFP and treated with Vehicle, BDNF, TrkB-Fc plus BDNF, or kn93 plus BDNF; n = 1649, 1742, 480, and 1380 trajectories, respectively (**h**), and in vehicle- or BDNF-treated neurons co-expressing FL-polyQ-HTT and GFP or GFP fused wild-type stargazin (Wt-stg-GFP), or ΔC stg, in which the binding domain to PSD95 was deleted; n = 495, 568, 376, 300, 573 and 498 trajectories, respectively (**i**). Diffusion coefficients were shown as median ± 25-75% IQR; significance was determined by unpaired two-tailed Student’s *t*-test (**b, d, e, f**), and Kruskal-Wallis test followed by Dunn’s Multiple Comparison Test (**g, h, i**); \**P* < 0.05, \*\**P* < 0.01, \*\*\**P* < 0.001.

We reasoned that if reduced CaMKII activity is responsible for aberrant AMPAR surface trafficking, then over-expression of constitutively active CaMKII should be able to rescue the FL-polyQ-HTT-induced increase in AMPAR surface diffusion. This is indeed what we observed (Fig. 3g). We assumed that if reduced CaMKII activity results from impaired BDNF-TrkB signaling pathway, then the application of exogenous BDNF should have similar effect. As expected, the application of exogenous BDNF similarly restored a lower GluA2-AMPAR surface diffusion (Fig. 3h, green bar). This rescue effect of BDNF requires the activation of TrkB and CaMKII as this effect was completely blocked by the addition of the BDNF scavenger TrkB-Fc or CaMKII inhibitor kn93 (Fig. 3h, orange and red bar, respectively). This indicates that BDNF-TrkB-CamKII signaling pathway plays a key role in stabilizing surface AMPARs.

The CaMKII-induced AMPAR immobilization requires the AMPAR auxiliary subunit stargazin and its binding to scaffold proteins of the postsynaptic density, such as PSD-95 ^32,36^. We thus examined the role of the interaction between stargazin and PSD-95 in mediating BDNF’s effects by expressing **Δ**C stargazin (**Δ**C Stg), in which the interaction domain with PSD-95 was deleted. In ΔC-Stg but not WT stargazin-expressing neurons, administration of BDNF failed to reduce GluA2-AMPAR surface diffusion (Fig.3i). These data suggest that impaired BDNF-TrkB-CaMKII signaling via the interaction between stargazin and PSD95 accounts for the disturbance of AMPAR surface diffusion in the hippocampus of HD models.

### Tianeptine improved BDNF protein production as well as intracellular trafficking in the hippocampus of HD models

BDNF is not a good candidate for HD treatment due to its instability and difficulties to cross the blood-brain barrier ^20,21,44^. An alternative approach, therefore, is to elevate endogenous BDNF protein or trafficking using other exogenous agents. Our previous work showed that the anti-depressant tianeptine modulates AMPAR surface diffusion and improved LTP in stress/depression models ^32^. It has also been reported that chronic tianeptine treatment increased BDNF protein level in various rodent brain structures ^45,46^. However, the effect of tianeptine on BDNF intracellular trafficking is not known and it is unclear whether tianeptine modulates BDNF signaling in HD models. We thus examined the effect of tianeptine on BDNF protein production as well as intracellular trafficking in different HD models. Hippocampal BDNF protein production was evaluated using ELISA and Western Blot methods in R6/1 and *Hdh*^Q111/Q111^ mice at 10-12 weeks of age. Because at this age, R6/1 and *Hdh*^Q111/Q111^mice were reported to show LTP defects and R6/1 mice gradually develop cognitive deficits ^4,6^. We found that the reduced hippocampal BDNF protein levels in R6/1 (Fig. 4a-c) and in *Hdh*^Q111/Q111^ mice (Fig. 4d) were both significantly improved by tianeptine administration (25mg/kg, i.p. daily for 4 days for R6/1 mice; and 10mg/kg, once for *Hdh*^Q111/Q111^ mice). Note that a single tianeptine injection at 10mg/kg was inefficient for R6/1 mice (data not shown), which may be due to the more severe phenotypes in this mouse model ^47^. We next examined tianeptine effect on BDNF intracellular trafficking in 3 different HD cellular models. The application of tianeptine fully rescued the velocity of BDNF anterograde and retrograde transport in polyQ-HTT-expressing neurons (Fig. 4e, 4f) as well as in hippocampal neurons from R6/1 (Fig. 4g) and *Hdh*^Q111/Q111^ mice (Fig.4h). In addition, tianeptine also augmented BDNF intracellular trafficking in WHTT-expressing neurons and in neurons from WT control for *Hdh*^Q111/Q111^ mice (Supple Fig. 3). These data suggest that tianeptine regulates hippocampal BDNF signaling at least at 2 levels, namely BDNF protein production and intracellular transport.

**Figure 4.**
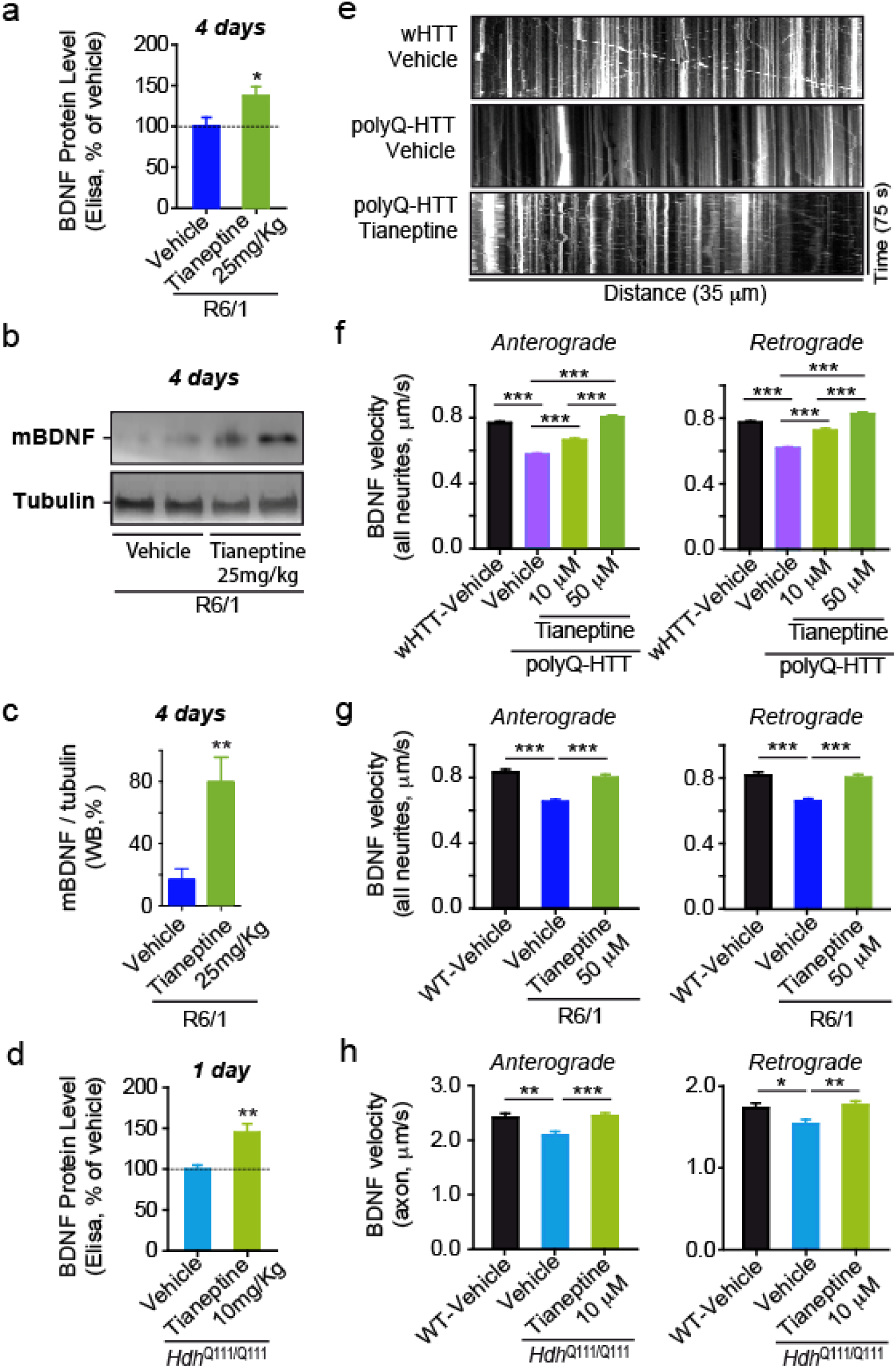
Antidepressant tianeptine rescued the reduced BDNF protein level and intracellular transport in different complementary HD models. (**a, b, c**) R6/1 mice were treated with saline (vehicle) or tianeptine (25 mg/kg, i.p. daily) for 4 days. Hippocampal BDNF protein level was assessed using ELISA Kit (**a**); values are mean ± s.e.m (% of vehicle); n = 14 and 13 mice for vehicle- and tianeptine-treated R6/1 group, respectively. Mature BDNF (mBDNF) and tubulin (for normalization) were analyzed by immunoblot (**b**); quantified densitometry of 14 KDa mBDNF, was expressed as percentage relative to tubulin (**c**); n = 9 and 7 mice for vehicle- and tianeptine-treated R6/1 group, respectively. (**d**) *Hdh*^Q111/Q111^ mice received one injection of saline or tianeptine (10mg/kg, i.p.). Hippocampal BDNF protein level was evaluated using ELISA kit; values are mean ± s.e.m (% of vehicle); n = 9 and 8 mice for vehicle- and tianeptine-treated *Hdh*^Q111/Q111^ group, respectively. (**e**) Representative kymographs of intracellular transport of BDNF-containing vesicles (white trajectories) in a neurite (35 μm from soma) over 75 seconds (s) in vehicle- or tianeptine-treated rat hippocampal neurons expressing wHTT or polyQ-HTT. (**f, g, h**) Anterograde and retrograde BDNF transport velocity in all neurites of vehicle- or tianeptine-treated wHTT- and polyQ-HTT-expressing rat hippocampal neurons (**f**), of hippocampal neurons from vehicle- or tianeptine-treated R6/1 mice and WT littermates (**g**), and in the axon of hippocampal neurons from vehicle- or tianeptine-treated *Hdh*^Q111/Q111^ and WT mice (**h**); values are mean ± s.e.m; n = 5569, 5656, 3339, 5737, 5227, 5706, 3190, and 5663 trajectories for anterograde and retrograde BDNF velocity in wHTT-vehicle, polyQ-HTT-vehicle, and polyQ-HTT-tianeptine (10 and 50 μM) neurons, respectively; n = 1424, 1710, 1512, 1376, 1487, and 1238 trajectories for anterograde and retrograde velocity in WT-vehicle, R6/1-vehicle, and R6/1-tianeptine (50 μM) neurons, respectively; n = 236, 261, 432, 194, 256, and 357 trajectories for anterograde and retrograde velocity in WT-vehicle, *Hdh*^Q111/Q111^ -vehicle, and *Hdh*^Q111/Q111^ -tianeptine (10 μM) neurons, respectively. Significance was determined by unpaired two-tailed Student’s *t*-test (**a, c, d**), and one-way ANOVA followed Bonferroni’s Multiple Comparison Test (**f, g, h**); \**P* < 0.05, \*\**P* < 0.01, \*\*\**P* < 0.001.

### >BDNF-TrkB signaling pathway mediates the tianeptine effect on BDNF intracellular trafficking and AMPAR surface diffusion

To further clarify the functional mechanism of tianeptine, we examined if the tianeptine-induced increase in BDNF intracellular trafficking could be prevented by a selective TrkB receptor inhibitor, Cyclotraxin-B (CB), which is a small inhibitor peptide mimicking the reverse turn structure of the variable region III that protrudes from the core of BDNF ^48^. Indeed, tianeptine (50 μM) induced improvement of anterograde and retrograde BDNF intracellular transport was fully blocked by pre-incubation with CB (1 μM) (Fig 5a, b). This suggested that tianeptine’s effect on BDNF intracellular trafficking is likely mediated through TrkB receptor. Since BDNF is not the sole ligand for TrkB receptor, we then postulated that if tianeptine influences BDNF intracellular trafficking through BDNF signaling rather than working in parallel, then addition of exogenous BDNF should be able to occlude tianeptine’s effect. Indeed, the administration of BDNF (100ng/ml) similarly rescued the decreased BDNF intracellular trafficking induced by polyQ-HTT and the combination of BDNF and tianeptine did not exhibit additive effect (Fig.5c). These data indicate that tianeptine affects BDNF intracellular trafficking possibly through BDNF-TrkB signaling pathway. We next asked whether tianeptine is also able to restore AMPAR surface traffic and if this effect is mediated by TrkB receptors. The application of tianeptine significantly slowed down AMPAR surface diffusion in polyQ-HTT-expressing neurons, an effect fully blocked by the TrkB receptor inhibitor Cyclotraxin-B and TrkB-Fc (Fig. 5 d, e). Collectively, these data suggest that the tianeptine effect on BDNF intracellular trafficking and AMPAR surface diffusion is mediated by BDNF-TrkB signaling pathway.

**Figure 5.**
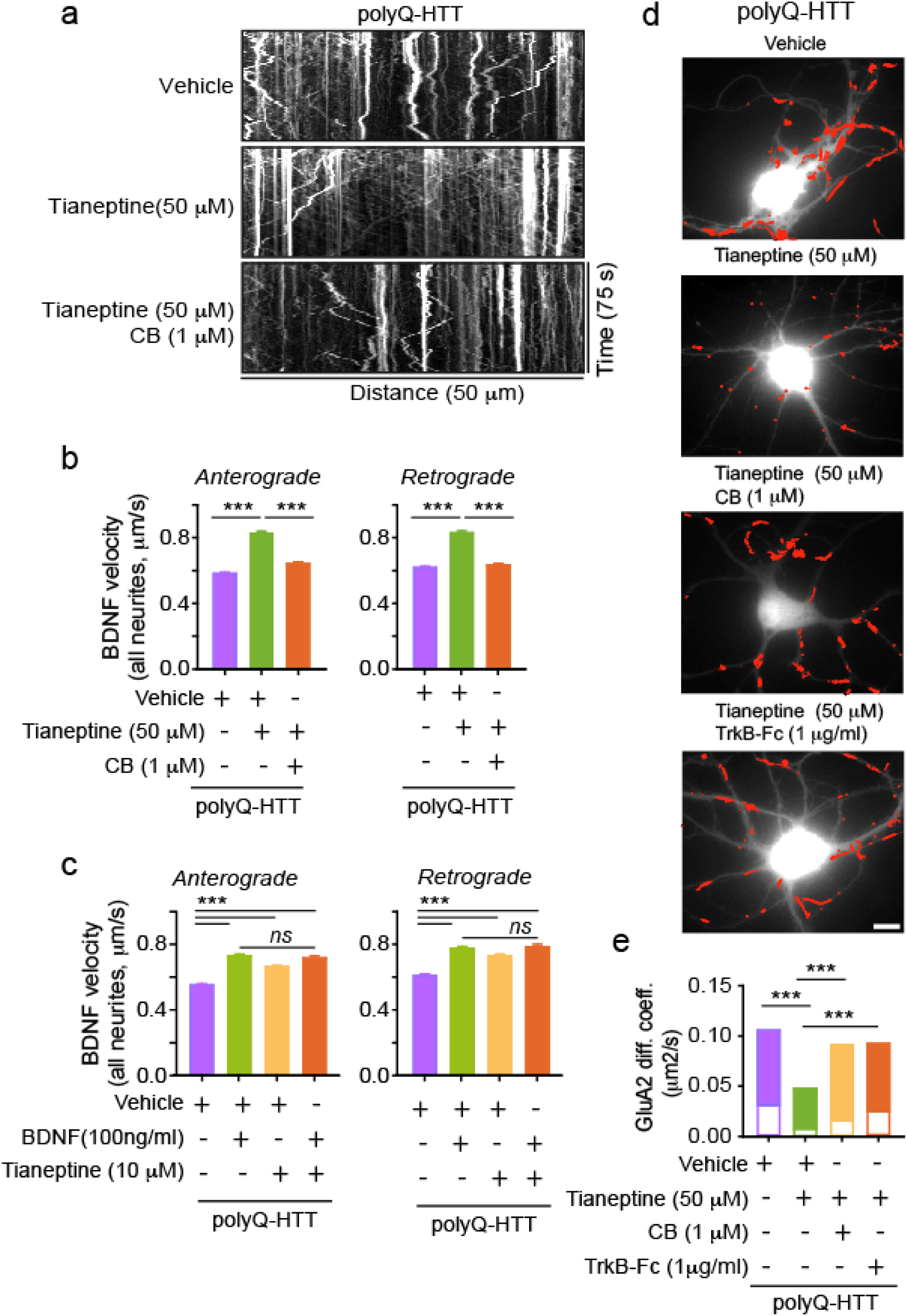
Tianeptine’s effect on BDNF intracellular transport and AMPAR surface diffusion is likely mediated by BDNF-TrkB signaling pathway. **(a)** Representative kymographs of intracellular transport of BDNF-containing vesicles (white trajectories) in a neurite (50 μm from soma) over 75 seconds (s) in polyQ-HTT-expressing rat hippocampal neurons treated with vehicle, tianeptine or cyclotraxin-B (CB) plus tianeptine. (**b, c**) Anterograde and retrograde BDNF transport velocity in all neurites of polyQ-HTT-expressing rat hippocampal neurons treated with vehicle, tianeptine or CB plus tianeptine (**b**), or treated with vehicle, BDNF, tianeptine, or BDNF plus tianeptine (**c**); values are mean ± s.e.m; n = 4322, 4017, 4199, 4354, 3887, and 3954 trajectories for anterograde and retrograde velocity in polyQ-HTT-expressing neurons treated with vehicle, tianeptine, and CB plus tianeptine, respectively (**b**); n = 3505, 3382, 3339, 2099, 3346, 3174, 3190 and 2022 trajectories for anterograde and retrograde velocity in polyQ-HTT-expressing neurons treated with vehicle, BDNF, tianeptine, and BDNF plus tianeptine, respectively (**c**). (**d**) Typical GluA2-QD trajectories (red) in polyQ-HTT-expressing rat hippocampal neurons, treated with vehicle, tianeptine, CB plus tianeptine, or TrkB-Fc plus tianeptine. Scale bars, 10μm. (**e**) GluA2-AMPAR diffusion coefficients in FL-polyQ-HTT-expressing rat hippocampal neurons treated with vehicle, tianeptine, CB plus tianeptine, or TrkB-Fc plus tianeptine; data are shown as median ± 25-75% IQR; n = 601, 535, 708, and 556 trajectories for 4 groups, respectively. Significance was assessed by one-way ANOVA followed Bonferroni’s Multiple Comparison Test (**b, c**) or Kruskal-Wallis test followed by Dunn’s Multiple Comparison Test (**e**); \**P* < 0.05, \*\**P* < 0.01, \*\*\**P* < 0.001; *ns*, not significant.

### Tianeptine enhanced the impaired hippocampal CA1 LTP and hippocampus-dependent memory as well as anxiety/depression-like behavior in complementary HD mouse models

BDNF-TrkB signaling and AMPAR surface diffusion are critically involved in hippocampal plasticity and learning and memory ^13,23^. We thus asked if tianeptine could rescue the impaired hippocampal LTP and hippocampal–dependent memory in 3 different mouse models of HD. Besides male heterozygous R6/1 transgenic mice and homozygous *Hdh*^Q111/Q111^ knock-in mice, we employed a 3rd mouse model, male CAG140 heterozygous knock-in mice, for behavior test. Heterozygous mice are highly relevant to the disease, as the majority of HD patients are heterozygous ^3^. In addition, CAG140 knock-in mice carry 140 polyQ and thus have earlier onset of symptoms than *Hdh*^Q111/Q111^ knock-in mice. The fEPSPs were recorded from mouse hippocampal CA1 neurons (Fig. 6a,b). R6/1 transgenic mice and *Hdh*^Q111/Q111^ knock-in mice were used at the age of 10-12 weeks. R6/1 mice showed a decrease in LTP of fEPSP slope compared to WT littermate control, which was partially rescued by chronic treatment of tianeptine at 25mg/kg (i.p. daily for 8 weeks) (Fig.6a) but not tianeptine at 10mg/kg (i.p. daily for 8 weeks)(data not shown). This suggested a dose-dependent effect of tianeptine. Very similar results were obtained in *Hdh*^Q111/Q111^ mice, in which LTP defects were normalized by a single injection of tianeptine (10mg/kg) (Fig. 6b). The restorative effect of tianeptine on LTP in HD mice raises the question of whether it can also rescue HD-related hippocampus-dependent cognitive impairments. In order to test early therapeutic intervention, we started to administer saline/tianeptine (10mg/kg, i.p. daily) to R6/1 and WT littermate mice from 4 weeks of age, when the mice do not typically present with cognitive deficits ^4^. At 12 weeks of age, the mice were subjected to open field test, Y-maze and contextual fear conditioning. The latter two tasks are hippocampal-dependent memory tasks ^49,50^, respectively based on novelty attractiveness and associated threat ^51^. Vehicle-treated R6/1 mice spent much less time in the novel arm than vehicle-treated WT mice, suggesting that R6/1 mice have impaired spatial working memory (Fig. 6c). Interestingly, tianeptine administration improved Y-maze performance of R6/1 mice, but not that of WT mice. This improvement is not due to a change in moving velocity, as vehicle- and tianeptine-treated R6/1 mice had similar moving velocity in open field (Supplemental Fig. 4a).

**Figure 6.**
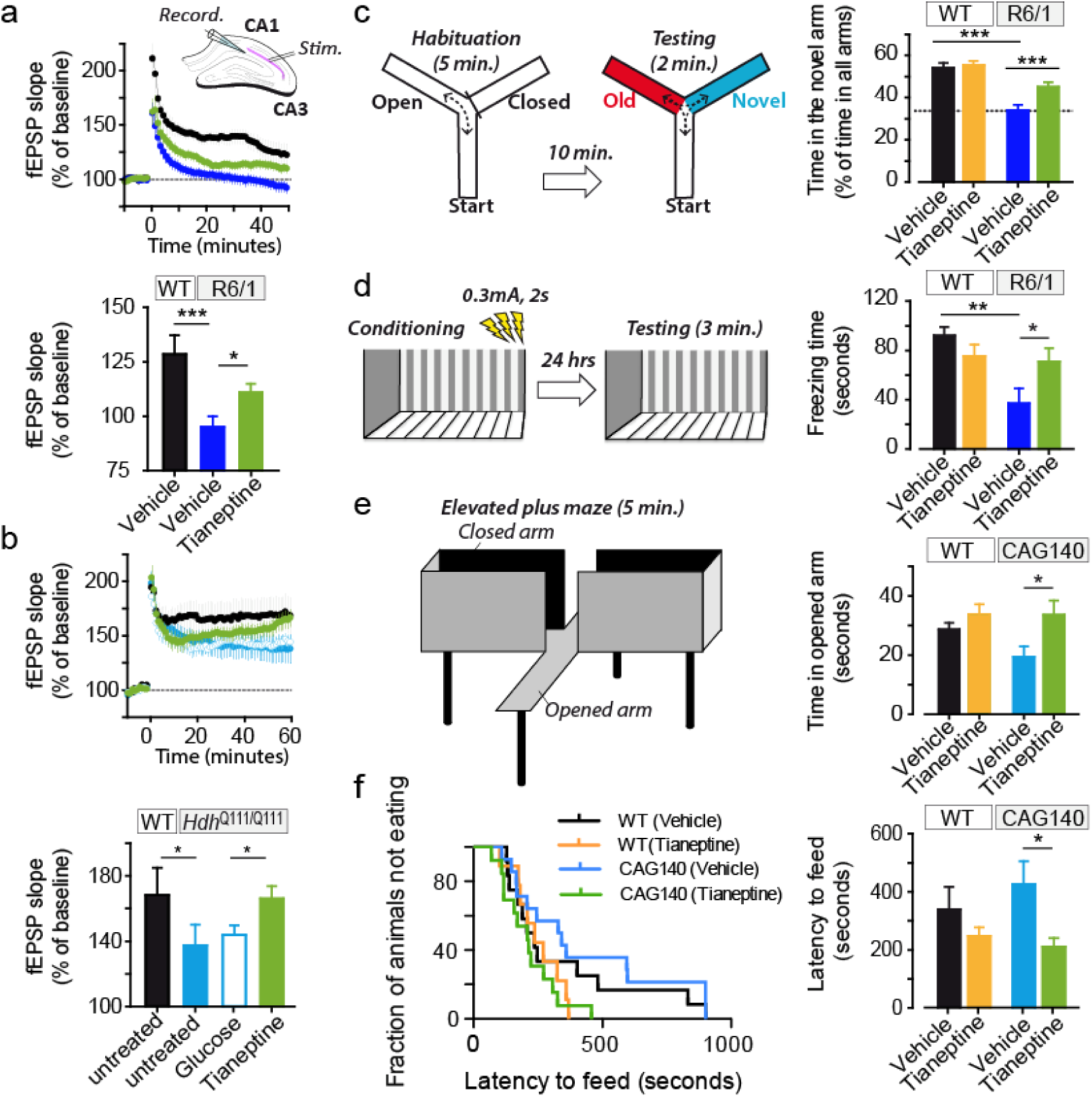
Tianeptine rescued impaired hippocampal CA1 LTP and hippocampus-dependent memory as well as anxiety/depression like behavior in different complementary HD mouse models. (**a, b**) Field EPSPs (fEPSPs) were recorded in CA1 region-containing acute slices of vehicle- or tianeptine-treated R6/1 |(**a**) and *Hdh*^Q111/Q111^ mice (**b**) following theta-burst stimulation of the Schaffer collaterals. Recording of fEPSPs was carried out blind with respect to genotype or treatment. Bar graph showing the percentage of potentiation observed during last 5-10 min of each recording; data are mean ± s.e.m; n = 16, 24, and 26 slices for vehicle-treated WT and R6/1 mice and tianeptine-treated R6/1 mice; n = 6, 11, 18, 16 slices for untreated WT, *Hdh*^Q111/Q111^ mice and glucose- and tianeptine-treated *Hdh*^Q111/Q111^ mice. (**c, d**) Hippocampus-dependent memory was examined using Y-maze (**c**) and contextual fear conditioning paradigm (**d**) in vehicle- or tianeptine-treated R6/1 and WT littermate mice. (**c**) Left, schematic diagram for Y-maze; right, percentage of time spent by mice in novel arms to that in total arms during 2-minute testing time. (**d**) Left, schematic diagram for contextual fear conditioning; right, freezing time during 3-minute testing time; data are mean ± s.e.m; n = 25, 28, 33, and 32 mice (**c**) and n = 10, 10, 10, and 12mice (**d**) for vehicle- and tianeptine-treated WT and R6/1 mice. (**e, f**) Anxiety/ depression-like behaviors were evaluated with elevated plus maze (EPM)(**e**) and novelty-suppressed feeding (NSF) paradigm (**f**) in HD CAG140 knock-in mice and WT littermates. (**e**) Left, schematic diagram for EPM; right, time spent in opened arms in EPM, which is an anxiety index. (**f**) Values plotted are cumulative survival of animals that did not eat over 15 minutes (left) or mean of latency to feed in seconds ± s.e.m (right). The latency to begin eating is an index of anxiety/depression-like behavior; n = 12, 9, 14, and 13 mice for vehicle- and tianeptine-treated WT and CAG140 mice (**e, f**). Significance was assessed by one-way ANOVA followed by Bonferroni’s Multiple Comparison Test (**a, b**), and two-way ANOVA followed by Bonferroni posttests (**c, d, e, f**). \**P* < 0.05, \*\**P* < 0.01, \*\*\**P* < 0.001.

Contextual fear conditioning, assessed by measuring the freezing behavior a mouse typically exhibits when re-exposed to a context in which a mild foot shock was beforehand delivered, reflects hippocampal-dependent memory ^49^. Vehicle-treated R6/1 mice exhibited less freezing in the contextual fear test compared to vehicle-treated WT littermates, indicating a worse memory, which was rescued by tianeptine treatment (Fig. 6d). Similarly, for the spatial working memory tested in Y-maze, no beneficial effect was observed on tianeptine-treated WT mice. We thus propose that tianeptine specifically rescued the hippocampal-dependent memory of R6/1 mice.

As HD mice also typically present an anxiety/depression-like phenotype ^47,52^, we asked if chronic tianeptine treatment would attenuate anxiety/depression-like phenotype in CAG140 heterozygous knock-in mice. The anxiety-like behavior of CAG140 mice was assessed using the Elevated Plus Maze (EPM) paradigm and Novelty Suppressed Feeding (NSF) paradigm. Anxiety-like phenotypes are characterized by decreased time spent in opened arms in EPM or an increase in latency to feed in NSF, which we observed in 6-month old CAG140 mice (data not shown). Here, we specifically investigated the early intervention therapy in HD and treated CAG140 mice mice starting from 3 months of age, when the anxiety phenotype is not fully established (compared to 6 months old mice, data not shown), and tested at 4 months of age. We found that compared to vehicle-treated CAG140 mice, chronically tianeptine-treated CAG140 mice spent significantly more time in opened arms in EPM (Fig. 6e), while their locomotor activity revealed by ambulatory distance was not significantly affected (Supplemental Fig.4b left panel). The treatment also markedly decreased the latency to feed in NSF (Fig. 6f) without altering the home food consumption (Supplemental Fig.4b right panel). In contrast, tianeptine did not significantly alter the behavior of WT mice, suggesting that chronic tianeptine treatment also specifically improves the anxiety/depression-like behavior in HD mice.

## DISCUSSION

AMPAR surface diffusion plays a key role in the regulation of the AMPAR synaptic content during glutamatergic synaptic plasticity ^22,30,31^. AMPARs constantly switch on the neuronal surface between mobile and immobile states driven by thermal agitation and reversible binding to stable elements such as scaffold or cytoskeletal anchoring slots or extracellular anchors. Even in synapses, AMPARs are not totally stable with around 50% of them moving constantly by Brownian diffusion within the plasma membrane, promoting continuous exchanges between synaptic and extrasynaptic sites^22^. This process is highly regulated by neuronal activity and other stimuli. It has been shown that the majority of AMPARs incorporated into synapses during LTP is from surface diffusion while exocytosed receptors likely serve to replenish the extrasynaptic pool available for subsequent bouts of plasticity ^31^. This AMPAR redistribution followed by immobilization and accumulation of AMPARs at synapses is the crucial step for the enhanced synaptic transmission during synaptic potentiation ^31,36,37^. In the present study, we provide the first direct proof in three complementary HD models that AMPAR surface mobility is significantly increased and that AMPARs fail to stabilize at the surface after cLTP stimuli. This opens a new perspective into the molecular mechanism underlying the impaired hippocampal synaptic plasticity in HD. It is noteworthy that disturbed AMPAR trafficking is also proposed to be one of the first manifestations of synaptic dysfunction that underlies Alzheimer’s Disease (AD) ^23,53,54^, which shares many clinical and pathological similarities with HD, such as early-onset cognitive deficiency before perceptible neuronal degeneration. Together with our previous finding that deregulated AMPAR surface diffusion underlies impaired LTP in stress/depression models^28^, these lines of evidence indicate that dysregulation of AMPAR surface diffusion may represent a common molecular basis for the impaired hippocampal synaptic plasticity and memory in various neuronal disorders.

It is generally accepted that BDNF via interaction with TrkB receptors enhances synaptic transmission and plasticity in adult synapses, while its binding to p75^NTR^ has been demonstrated to negatively modulate synaptic plasticity, spine-dendrite morphology and complexity ^13^. Recent studies show that impaired BDNF delivery, as well as the abnormally reduced expression of TrkB receptor and enhanced p75^NTR^ expression account for the hippocampal synaptic and memory dysfunction ^9,18,19^. The phosphorylation of GluA1 on Ser-831 through activation of protein kinase C and CaMKII via TrkB receptors has been proposed to be responsible for AMPAR synaptic delivery ^13^. However, other evidence indicates that GluA1 phosphorylation at Ser-831 alters single-channel conductance rather than receptor anchoring ^55^. Here we provide the first evidence in HD models that administration of BDNF slows down the increased AMPAR surface diffusion via interaction between PSD95 and stargazin, which is downstream of TrkB-CaMKII signaling pathway. It is possible that both processes, that is, change in the single-channel conductance and receptor anchoring (this study), occur in parallel and affect AMPAR signaling. Interestingly, the reduced CaMKII activity reported in *Hdh*^Q111/Q111^ knock-in mice could be prevented by normalization of p75^NTR^ levels ^9^. This effect could be attributable to the preservation of TrkB signaling, as it has been shown that decreasing p75 ^NTR^ expression or blocking its coupling to the small GTPase RhoA normalizes TrkB signaling; while upregulation of p75 ^NTR^ signaling through phosphatase-and-tensin-homolog-deletedon-chromosome-10 (PTEN) results in impaired TrkB signaling ^19^. Thus, impaired BDNF delivery and aberrant processing of BDNF signal may converge on TrkB-CaMKII signaling pathway affecting AMPAR surface diffusion.

Tianeptine is a well-tolerated antidepressant primarily used in the treatment of major depressive disorders ^56^. It is structurally similar to a tricyclic antidepressant (TCA), but has different pharmacological properties than typical TCAs as it produces its antidepressant effects likely through the alteration of glutamate receptor activity ^32,56^. Tianeptine alters glutamatergic transmission, increasing for instance the phosphorylation of GluA1 subunits ^57^ and activating CaMKII and protein kinase A via the p38, p42/44 mitogen-activated protein kinases (MAPK) and c-Jun N-terminal kinases (JNK) pathways ^58^. Through unknown mechanisms, tianeptine prevents stress-induced dendritic atrophy, improves neurogenesis, reduces apoptosis and normalizes metabolite levels and hippocampal volume ^56^. In the present study, we show in complementary HD models that tianeptine restored AMPAR surface diffusion, via BDNF-TrKB signaling pathway, and rescued defective LTP and hippocampal-dependent memory. Interestingly, the activation of BDNF-TrkB signaling pathway is also required for the effect on the depression-like behavior of some typical antidepressants, such as fluoxetine and imipramine ^59,60^. The mechanisms underlying chronic tianeptine treatment may involve BDNF-induced neurogenesis ^56^, however, our finding that a single dose administration of tianeptine is sufficient to rescue aberrant LTP in *Hdh*^Q111/Q111^ knock-in mice points to additional mechanisms. Given the critical role of AMPAR surface diffusion in hippocampal synaptic plasticity ^23,31,36,37^, we argue that the beneficial effects of tianeptine on the impaired LTP and hippocampal-dependent memory stem, at least in part, from its normalization of AMPAR surface diffusion. Although tianeptine is also able to augment BDNF intracellular trafficking in WT controls (Supple Fig. 3) and immobilize AMPAR surface diffusion under basal conditions ^32^, it did not significantly improve hippocampal-dependent memory in WT mice, suggesting that the maintenance of a physiological dynamic equilibrium is key to an effective treatment. The present study also showed beneficial effect of tianeptine on the anxiety/depression-like behavior in CAG140 knock-in mouse model. Note that cognitive dysfunction and psychiatric pathologies such as depression, stress and anxiety are typical features of HD, which occur well before the onset of motor dysfunction ^1,2,34,61,62^, thus the use of tianeptine may represent a promising early therapeutic strategy for HD targeting both psychiatric and cognitive defects. Moreover, that tianeptine is a clinically used drug will also facilitate clinical trials.

In conclusion, we unravel AMPAR surface diffusion as a potential novel therapeutic target for early intervention in HD and propose a new therapeutic strategy for HD using an antidepressant tianeptine, which improved hippocampal synaptic and memory deficits as well as anxiety/depression-like behavior in HD mice possibly through the modulation of BDNF signaling and AMPAR surface diffusion.

## MATERIALS AND METHODS

### HD transgenic mice, primary Neuronal Cultures and transfection

The heterozygous male R6/1 mice (Jackson Laboratory, Main Harbor, NY) were crossed with female C57BL/6 mice (Charles River, Lyon). Homozygous *Hdh*^Q111/Q111^ KI mice of HD on CD1 background are generous gift from M.E. MacDonald ^63^. The CAG140 are heterozygous mice with C57Bl6N/J background. The animals were housed with food and water ad libitum under a 12h light–dark cycle. All work involving animals was conducted according to the rules of ethics of the Committee of University of Bordeaux and the Aquitaine (France) and the Institutional Animal Care and Use Committee (European Directive, 2010/63/EU for the protection of laboratory animals, permissions # 92-256B, authorization ethical committee CEEA 26 2012_100). Polymerase chain reaction (PCR) genotyping with DNA extracted from a piece of tail was carried to identify mice genotype.

Primary cultures of hippocampal neurons were prepared following a previously described method from (1) Sprague-Dawley rats at E18; (2) *Hdh*^Q111/Q111^ KI mice and WT littermates at P0 for AMPAR surface tracking and *Hdh*^Q111/Q111^ KI mice and WT mice at E15 for BDNF intracellular tracking; (3) *R6*/1 mice and WT littermates at P0 ^10,64^. Cells were plated at a density of 200 × 10^3^ cells for rat culture and 450 × 10^3^ cells for mice culture per 60 mm dish on poly-lysine pre-coated cover slips. Cultures were maintained in serum-free neurobasal medium (Invitrogen) and kept at 37 °C in 5% CO2 for 20 div at maximum. Cells were transfected with appropriate plasmids using Effectene (Qiagen).

### Plasmid Constructs & Chemical product

Homer 1C::GFP with CaMKII promoter was generated by subcloning homer 1C cDNA into the eukaryotic expression vector pcDNA3 (Invitrogen); EGFP was inserted at the N-terminus of the Homer 1C sequence. Exon1 mutant huntingtin contains 69 polyglutamine expansion (exon1-polyQ-HTT) and wild-type huntingtin with 17 polyglutamine (exon1-wHTT) ^35^. Full-length HTT plasmids encode full-length huntingtin with 17 polyQ (FL-wHTT) or 75Q (FL-polyQ-HTT). 480-17Q, 480-68Q huntingtin plasmids encode the first 480 amino acids fragment of huntingtin with 17 (Nter-wHTT) or 68 glutamines (Nter-polyQ-HTT) ^65,66^. Tianeptine was purchased from T & W group and MedChemexpress CO.,Ltd; BDNF from Sigma-Aldrich; TrkB-Fc from R&D Systems; kn93 from Tocris. Homemade Cyclotraxin-B and Cyclotraxin-B synthesized by Bio S&T were used.

### Cyclotraxin B synthesis

Cyclotraxin B was synthesized at a 0.05 mmol scale. Amino acids were assembled by automated microwave solid phase peptide synthesis on a CEM microwave–assisted Liberty-1 synthesizer following the standard coupling protocols provided by the manufacturer. Methionine was replaced by Norleucine (λ), a more stable isostere. Linear peptide was cleaved (TFA:H2O:EDT:TIS, 94:2.5:2.5:1) and purified by HPLC. Disulfide bond formation was carried out for 10 hrs in H20 in the presence of DMSO (5%) and ammonium acetate (0.05 M) at high dilution of the peptide (100 µM). Solvent excess was removed and the peptide was purified by RP-HPLC (YMC C18, ODS-A 5/120, 250×20 mm, UV detection at 228 and 280 nm, using a standard gradient: 5% MeCN containing 0.1% TFA for 5 min followed by a gradient from 10 to 40% over 40 min in dH2O containing 0.1% TFA at a flow rate of 12 mL.min-1). Peptides were characterized by analytic RP-HPLC and MALDI. Peptides were quantified by absorbance measurement at 280 nm, aliquoted, lyophilized and stored at -80 °C until usage.

### Single nanoparticle (Quantum dot) tracking and surface diffusion calculation

Rat primary hippocampal neurons were co-transfected at DIV 10-11 with GFP/homer1c-GFP and wHTT/polyQ-HTT at the ratio of 1:9 to ensure that the majority of GFP-transfected neurons were transfected with HTT. Homer1c was used as a postsynaptic marker. Endogenous GluA2 and GluA1 quantum dot (QD) tracking was performed at DIV 11-12 as previously described ^32^. Neurons were first incubated with mouse monoclonal antibody against N-terminal extracellular domain GluA2 subunit (a kind gift from E. Gouaux, Oregon Health and Science Unieristy, USA) or rabbit poloclonal antibody against N-terminal extracellular domain GluA1 subunit (PC246, Calbiochem) followed incubation with QD 655 Goat F(ab’)2 anti-mouse or anti-Rabbit IgG (Invitrogen). Non-specific binding was blocked by 5% BSA (Sigma-Aldrich). QDs were detected by using a mercury lamp and appropriate excitation/emission filters. Images were obtained with an interval of 50 ms and up to 1000 consecutive frames. Signals were detected using a CCD camera (Quantem, Roper Scientific). QDs were followed on randomly selected dendritic regions for up to 20 min. QD recording sessions were processed with the Metamorph software (Universal Imaging Corp). The instantaneous diffusion coefficient, D, was calculated for each trajectory, from linear fits of the first 4 points of the mean-square-displacement versus time function using MSD(t) = <r^2^> (t) = 4Dt. The two-dimensional trajectories of single molecules in the plane of focus were constructed by correlation analysis between consecutive images using a Vogel algorithm. QD-based trajectories were considered synaptic if colocalized with Homer 1C dendritic clusters for at least five frames.

### BDNF intracellular transport

Rat hippocampal neurons were co-transfected at DIV 9-10 with GFP-fused 480-17Q (GFP::Nter-wHTT) or 480-68Q (GFP::Nter-polyQ-HTT) and mCherry-BDNF at the ratio of 4:1 using Effectene (QIAGEN). Live imaging was carried out at DIV 10-11. The movement of BDNF-containing vesicles was tracked using video-microscopy on an inverted Leica DMI 6000 Year microscope (Leica Microsystems, Wetzlar, Germany) equipped with a HQ2 camera (Photometrics, Tucson, USA). The objective HCX PL used was a CS APO 63X NA 1.32 oil. The atmosphere was 37 °C incubator created with year box and air heating system (Life Imaging Services, Basel, Switzerland). Acquisitions and calculation were done on the MetaMorph software (Molecular Devices, Sunnyvale, USA). For BDNF axonal trafficking in hippocampal neurons of mouse *Hdh*^Q111/Q111^ KI and WT mice, hippocampal neurons at E15 were used. Microchambers, neuronal transfection as well as videomicroscopy were previously described ^42^. Images were collected in stream mode using a Micromax camera (Roper Scientific) with an exposure time of 100 to 150 ms. Projections, animations and analyses were generated using ImageJ software (http://rsb.info.nih.gov/ij/, NIH, USA). Maximal projection was performed to identify the vesicles paths, which in our system corresponds to vesicle movements in axons. Kymographs and analyses were generated with the KymoToolBox, a home-made plug-in ^42^.

## UPAINT

Rat primary hippocampal neurons were co-transfected at DIV 4 with homer1c-GFP and FL-wHTT/polyQ-HTT for 2 weeks. Homer1c was used as a postsynaptic marker. Single-molecule fluorescent spots were localized in each frame and tracked over time as previously described ^39^.

### BDNF enzyme-linked immunosorbent assay (ELISA)

The BDNF concentration was evaluated using BDNF ELISA Kit (Millipore, Abnova).

### Fluorescence Resonance Energy Transfer (FRET)- Fluorescence-lifetime imaging microscopy (FLIM) experiments

A FRET-based CamKIIα, named REACh-CamKII is a kind gift from R. Yasuda (Max Planck Insitute, Florida, USA). The amino and carboxy termini of CamKIIa are labeled with the FRET pair of monomeric enhanced green fluorescent protein (mEGFP) and resonance energy-accepting chromoprotein (REACh), a non-radiative yellow fluorescent protein variant. ^43,67^. FLIM experiments were performed at 37 °C using an incubator box with an air heater system (Life Imaging Services) installed on an inverted Leica DMI6000B (Leica Microsystem) spinning disk microscope and using the LIFA frequency domain lifetime attachment (Lambert Instruments, Roden, The Netherlands) and the LI-FLIM software. Cells were imaged with an HCX PL Apo X 100 oil NA 1.4 objective using an appropriate GFP filter set. Cells were excited using a sinusoidally modulated 1-W 477nm LED (lightemitting diode) at 40 MHz under wild-field illumination. Emission was collected using an intensified CCD LI2CAM camera (FAICM; Lambert Instruments). The phase and modulation were determined from a set of 12 phase settings using the manufacturer’s LI-FLIM software. Lifetimes were referenced to a 1µM solution of fluorescein in in Tris-HCl (pH 10) that was set at 4.00 ns lifetime. Signals were recorded with a back-illuminated Evolve EMCCD camera (Photometrics). Acquisitions were carried out on the software MetaMorph (Molecular Devices).

### Western blotting

Western Blot is performed as previously described ^32^. 10 µg of protein was loaded per lane and analyzed by SDS–PAGE. Primary antibodies anti-BDNF antibody (Santa Cruz Biotechnology, sc-546); anti-Tubulin antibody (Sigma-Aldrich) were used.

### Ex vivo extracellular recording from hippocampal CA1 pyramidal neurons

Male heterozygous R6/1 mice, *Hdh*^Q111/Q111^ KI mice and respective WT littermates were used for ex vivo extracellular recording. *Hdh*^Q111/Q111^ KI mice (10-12 week of age) received a single injection of tianeptine (i.p., 10mg/kg) with saline as negative control; R6/1 mice received chronic tianeptine treatment (25mg/kg, i.p. daily) starting from 4 weeks of age until 12 weeks of age. As described previously ^32^, a hippocampal slice was transferred to a superfusing recording chamber with temperature controlled at 33.5 °C, and continuously perfused with oxygenated ACSF using a peristaltic pump (Ismatec, Switzerland). A teflon-coated tungsten bipolar stimulating electrode (Phymep, Paris, France) was positioned in stratum radiatum, allowing the afferent schaffer collateral-commissural pathway from the CA3 area to the CA1 region to be stimulated. The field-EPSPs (fEPSPs) were recorded from stratum radiatum of CA1 area, using a glass electrode (3–5 MΩ) pulled from borosilicate glass tubing (Havard Apparatus, USA; 1.5 mm O.D x 1.17 mm I.D) and filled with ACSF. Pulses were delivered at 7.5s by a stimulus isolator (Isoflex, AMPI, Jerusalem, Israel), with adjusting current intensity to obtain 30-40 % of the maximum fEPSP. A theta-burst stimulatio (TBS) protocol (4 pulses, respectively, delivered at 100 Hz, repeated 10 times, at an interval of 200 ms) was delivered by Clampex10.4 (Molecular Devices, USA) and the stimulus isolator to induce LTP. Recordings were made continually for more than 60 min, following the TBS. Data were recorded with a Multiclamp700B (Axon Instruments, USA) and acquired with Clampex10.4. The slope of the fEPSP was measured using clampfit10.4 software, with all values normalized to a 5 min baseline period; the values during 50-60 min after TBS are reported in the figures as ± standard error of the mean (SEM). Mean values were compared between genotypes and treatments using either unpaired Student’s t-test as appropriate. The experiments were done blindly.

### Behavioral tests

Male R6/1 and WT littermate mice were used for behavioral tests. At 4 weeks of age, littermate mice with mixed genotypes were housed (3-5 per cage) in polycarbonate standard cages (33×15×14cm) and randomly allocated to vehicle or drug treatment groups. Mice received daily intraperitoneal injection (i.p.) of 0.9% saline (vehicle) or tianeptine (10mg/kg) dissolved in 0.9% saline until 12 weeks of age, when the animals were subjected to a battery of behavioral tests. On day 1, all mice were subjected to Open Field test; on day 2, spatial memory was assessed in Y maze. Following a week rest, on day 9, a subset of mice were further tested for contextual fear conditioning, which is performed lastly in order to minimize confounding factors. All behavioral testing was carried out in the light phase (light intensity: 45-50 lux). Before each behavioral test, mice were individually housed in standard cages with sawdust, food and water and left undisturbed in the experimental room at least 30min before testing began.

### Open field

The apparatus constituted of a white square arena (42cm x 42cm x 20 cm). Each animal was placed in the center of the arena and allowed to explore for 20 min. Images tracked from a camera above the maze were analyzed with Ethovision (version 9.1). The total distance traveled and the time spent moving were analyzed as readouts of locomotor activity. The apparatus was cleaned by ethanol 70% between mice.

### Y-maze

Hippocampal-dependent spatial working memory was evaluated using Y-maze. The apparatus consisted of three identical grey plastic arms (42 × 8 × 15 cm) and spaced at 120° of each other. The maze was located in the middle of a room containing a variety of extramaze cues. A digital camera was mounted above the maze transmitting the data to a PC running the Ethovision system. Mice were assigned two arms (start and familiar arm) to which they were exposed during the first phase of the test (sample phase). The remaining third arm blocked by a gray plastic door constituted the novel arm during the second phase (test phase). Mice were placed at the end of the start arm and allowed to explore freely both the start and the other unblocked arm for 5 min before being removed from the maze and returned to the waiting cage. After 10 min in the waiting cage, the test phase began. During this phase, the door was removed and all three arms were unblocked; mice were placed at the end of the start arm and allowed to explore the entire maze for 2 min. Timing of both the sample and test phase periods began once the mouse had left the start arm. The apparatus was cleaned between the two phases in order to avoid olfactory cues. Time spent in the novel arm in comparison to time in all three arms was used as one readout for hippocampal-dependent spatial memory.

### Contextual fear conditioning

Contextual fear conditioning provides a measure of memory by assessing a memory for the association between mild foot shock and a salient environmental cue. In the fear conditioning test, freezing behavior is defined as the complete lack of movement, which is a characteristic fear response in rodents, providing a readout of hippocampal-dependent memory. Fear conditioning was performed in a testing chamber with internal dimensions of 25 × 25 × 25 cm, which has transparent plastic walls each side and steel bars on the floor. A camera mounted at one side recorded each session. The chamber was located inside a larger, insulated, transparent plastic cabinet (67 × 53 × 55 cm) that provided protection from outside noise. The cabinet contained a ventilation fan that was operated during the sessions. Mice were held outside the experimental room in individual cages prior to testing. Training chambers were cleaned with 100% ethanol solution before and after each trial to avoid any olfactory cues. The experiments ran over two consecutive days. On Day 1, mice were placed in the conditioning chamber and 2 min 28s later received one footstock (2 s, 0.3 mA). Mice were removed from the chamber 30 s after the shock. On Day 2, they returned to the same conditioning chamber for a 3-min period in the exact same conditions as Day 1, but without electrical shock, to evaluate context-induced freezing.

### Elevated plus maze (EPM)

Male CAG140 heterozygous knock-in mice received daily i.p. injection of saline or tianeptine at 10mg/kg at 12 weeks of age. Behavioral tests were performed at 16 week of age. Each animal, over a week, was successively tested in the Elevated Plus Maze (EPM) and Novelty Suppressed Feeding (NSF), which represent different anxiety and depression behavior paradigms. Behavioral tests were performed during the light phase between 0700 and 1900. EPM was performed as previously ^68^. The maze is a plus-cross-shaped apparatus, with two open arms and two arms closed by walls linked by a central platform 50 cm above the floor. Mice were individually put in the center of the maze facing an open arm and were allowed to explore the maze during 5 min. The time spent in and the number of entries into the open arms were used as an anxiety index. Locomotion was also measured to ensure any confounding effects. All parameters were measured using a videotracker (EPM3C, Bioseb, Vitrolles, France).

### Novelty-Suppressed-Feeding (NSF)

The NSF is a conflict test that elicits competing motivations: the drive to eat and the fear of venturing into the center of a brightly lit arena. The latency to begin to eat is used as an index of anxiety/depression-like behavior, because classical anxiolytic drugs as well as chronic antidepressants decrease this measure. The NSF test was carried out during a 15-min period as previously described ^68^. Briefly, the testing apparatus consisted of a plastic box (50×50×20 cm), the floor of which was covered with approximately 2 cm of wooden bedding. Twenty-four hours prior to behavioral testing, all food was removed from the home cage. At the time of testing, a single pellet of food (regular chow) was placed on a white paper platform positioned in the center of the box. Each animal was placed in a corner of the box, and a stopwatch was immediately started. The latency to eat (defined as the mouse sitting on its haunches and biting the pellet with the use of forepaws) was timed. Immediately afterwards, the animal was transferred to its home cage, and the amount of food consumed by the mouse in the subsequent 5 min was measured, serving as a control for change in appetite as a possible confounding factor.

### Statistics

For imaging data, statistical values are given as mean ± SEM. or medians ± interquartile range (IQR) defined as the interval between 25% - 75% percentile. Statistical significances were tested using Prism 6.0 (GraphPad, USA). Normally distributed data sets were compared using the paired or unpaired Student’s *t*-test. Statistical significance between more than two normally distributed datasets was tested by one way ANOVA variance test followed by a Bonferroni test to compare individual pairs of data. Non-Gaussian data sets were tested by non-parametric Mann-Whitney test. For behavioral tests, statistical analysis was carried out by two-way ANNOVA with genotype and treatment as the between-subject factors. Indications of significance correspond to *p* values < 0.05 (*), *p* < 0.01 (**) and *p* < 0.001 (***).

## ACKNOWLEDGMENTS

We acknowledge E. Gouaux for the anti-GluA2 antibody; J.B. Sibarita for providing single particle analysis software; C Poujol, S Marais, F Cordelieres from Bordeaux Imaging Center, part of the France BioImaging national infrastructure, for support in microscopy; and C. Breillat, E. Verdier, N Retailleau for cell culture and plasmid production. The guidance of Françoise Coussen for biochemistry is acknowledged. We thank Jia-Yi Li for valuable input in the early phase of this project. This work was supported by funding from the Conseil Régional d’Aquitaine, ANR NanoDom and Stim-Traf-Park, Labex BRAIN and ANR-10-INBS-04 France-BioImaging, Centre National de la Recherche Scientifique, ERC grant nanodyn-syn and ADOS to D.C. and Aquitaine Science Transfer grant to D.C and H.Z. and FRM to H.Z.

## >AUTHOR CONTRIBUTIONS

H.Z. performed AMPAR and BDNF trafficking as well as live cell imaging studies, C.Z. performed electrophysiological experiments, J.V. conducted biochemical studies and behavioral test in R6/1 mouse line and assisted in imaging analysis, D.Z. performed BDNF trafficking studies in *Hdh*^Q111/Q111^ mouse neurons, D.J.D and C.B. conducted behavioral test in CAG140 mouse line; D.J.D also contributes to the interpretation of behavioral tests in CAG140 mouse line; M.S. and D.G. synthesized Cyclotraxin-B; Y.C contributed to supervision and interpretation of behavioral studies in R6/1 mouse line, F.S contributed to the supervision and interpretation of BDNF trafficking studies in *Hdh*^Q111/Q111^ mouse neurons and behavioral tests in CAG140 mouse line, Y.H. contributed to the supervision of electrophysiology studies and format of all figures. D.C. and H.Z. developed the concept, supervised the project, contributed to the design and interpretation of all experiments, and wrote the manuscript. All authors had the opportunity to discuss results and comment on the manuscript.

---

**Supplemental Figure 1.**
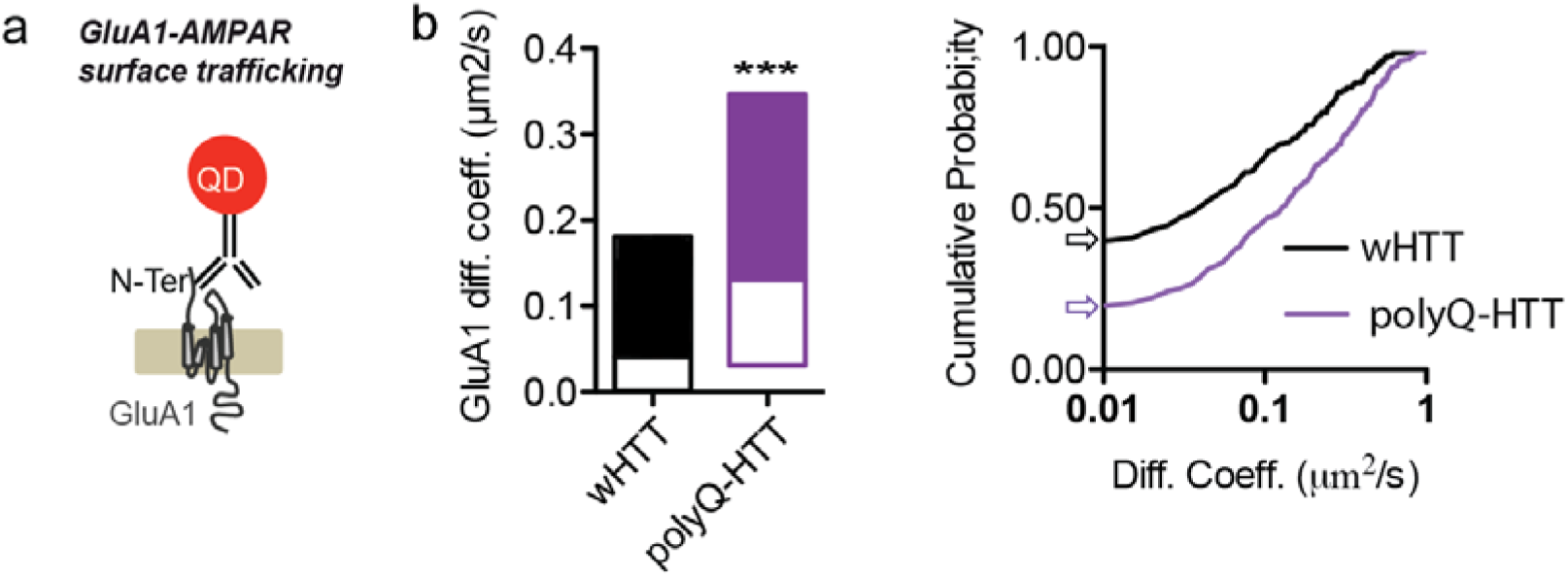
Deregulated GluA1-AMPAR surface diffusion in rat hippocampal neurons expressing FL-polyQ-HTT **(a)** Experimental scheme showing that for endogenous GluA1-AMPAR surface tracking, hippocampal neurons were incubated with rabbit polyclonal antibody against N-terminal extracellular domain of GluA1 subunit followed by QD anti-rabbit IgG. (**b**) Left, GluA1-AMPAR diffusion coefficients in rat hippocampal neurons expressing FL-wHTT or FL-polyQ-HTT; data are shown as median ± 25-75% IQR; n = 206 and 310 trajectories, respectively. right, cumulative probability of GluA1 diffusion coefficient. The first point of the probability corresponding to the fraction of immobile receptors with diffusion coefficients ≤ 0.01 μm2/s was showed by arrows. The cumulative curve of FL-polyQ-HTT expressing neurons shifts toward right implying an increased GluA1-AMPAR surface diffusion. Significance was determined by Mann-Whitney test. \*\*\**P*< 0.001.

**Supplemental Figure 2.**
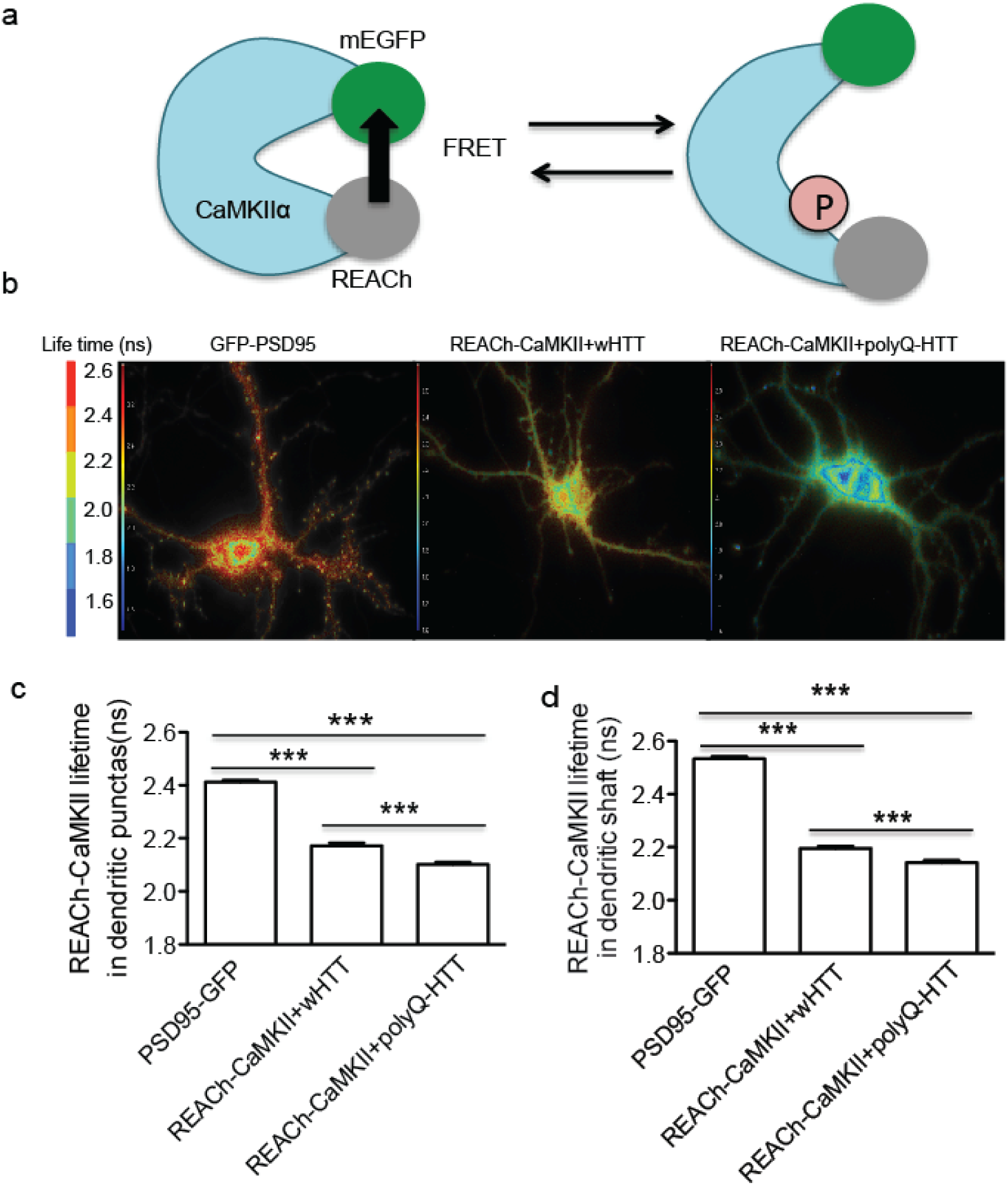
Decreased CaMKII activity in HD cellular model (**a**) Schematic diagram showing fluorescence resonance energy transfer (FRET)-based CamKIIα, named REACh-CamKIIα. The activation of REACh-CaMKII changes the conformation to the open state in which its kinase domain is exposed, thereby decreasing FRET and increasing the fluorescence lifetime of mEGFP. (**b**) Representative lifetime image of rat hippocampal neurons expressing PDS95-GFP, REACh-CaMKIIα plus FL-wHTT, and REACh-CaMKIIα plus FL-polyQ-HTT. Blue color indicates strong FRET and short lifetime, while red color represents weak FRET and long lifetime. (**c, d**) Quantification of lifetime in randomly-selected regions in dendritic puncta (**c**) or in dendritic shaft (**d**) in rat hippocampal neurons expressing PDS95-GFP, REACh-CaMKIIα plus FL-wHTT, or REACh-CaMKIIα plus FL-polyQ-HTT. PDS95-GFP-expressing neurons showed long lifetime (≥ 2.4 ns) in both dendritic puncta and shaft indicating no FRET. Lower lifetime indicates stronger FRET and reduced CamKIIα activity; data are mean ± s.e.m; n = 178, 231, and 238 regions for dendritic puncta, and n = 93, 115 and 188 regions for dendritic shaft in neurons expressing PDS95-GFP, REACh-CaMKIIα plus FL-wHTT, and REACh-CaMKIIα plus FL-polyQ-HTT, respectively. Significance was assessed by One-way ANOVA followed by Bonferroni’s Multiple Comparison Test; *** *P* < 0.001.

**Supplemental Figure 3.**
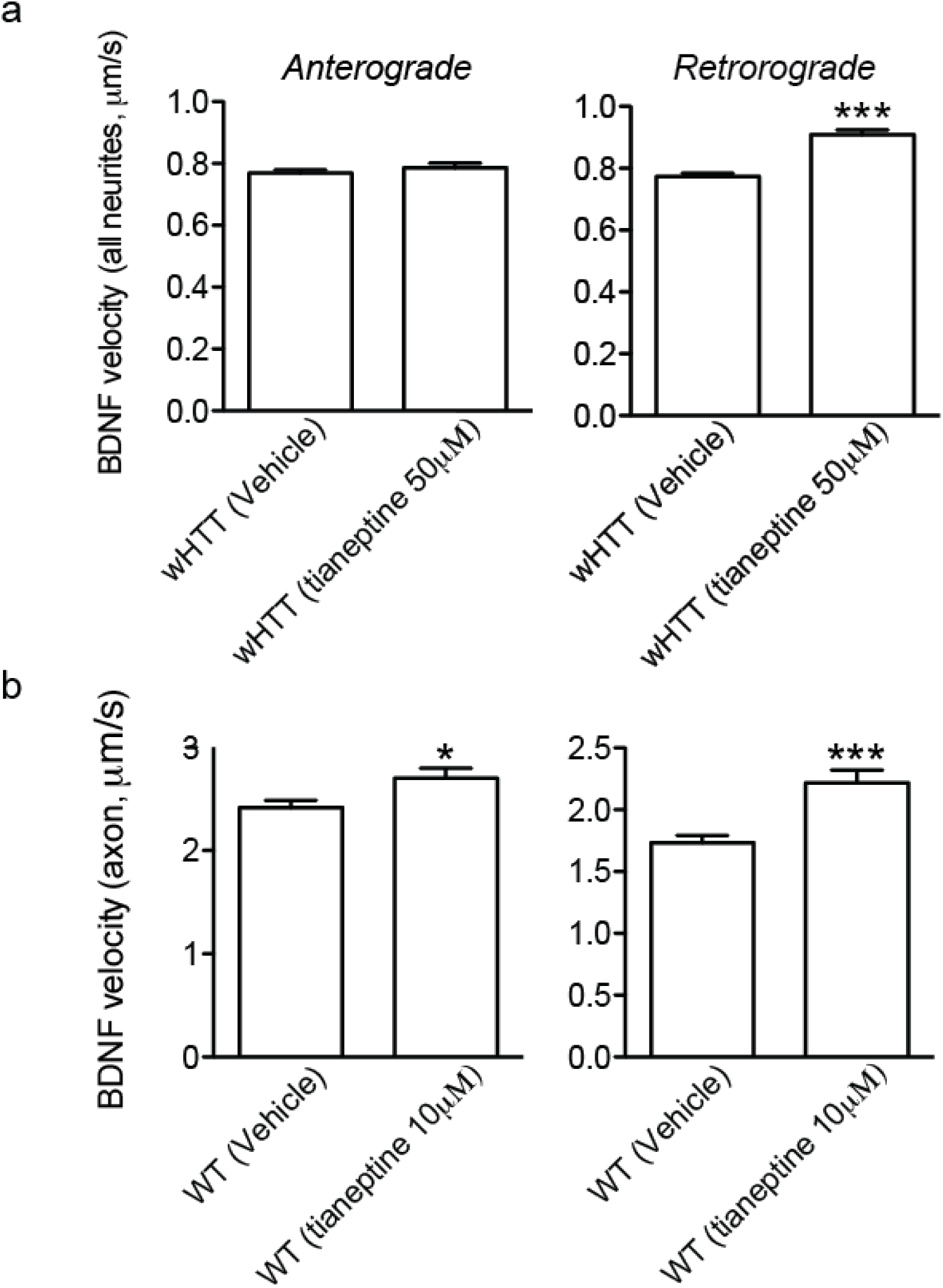
Tianeptine facilitated BDNF intracellular transport in wHTT-expressing rat hippocampal neurons and neurons from WT mice. (**a, b**) Anterograde and retrograde BDNF transport velocity in all neurites of vehicle- or tianeptine-treated wHTT-expressing rat hippocampal neurons (**a**), and in the axon of hippocampal neurons from WT mice for *Hdh*^Q111/Q111^ mouse line (**b**); values are mean ± s.e.m; n = 5569, 2522, 5227 and 2542 trajectories for anterograde and retrograde BDNF velocity in vehicle and tianeptine-treated wHTT-expressing neurons, respectively; n = 236, 157, 194 and 110 trajectories for anterograde and retrograde BDNF velocity in vehicle and tianeptine-treated neurons from WT mice for *Hdh*^Q111/Q111^ mouse line. Significance was determined by unpaired two-tailed Student’s *t*-test; \**P* < 0.05, \*\*\**P* < 0.001.

**Supplemental Figure 4.**
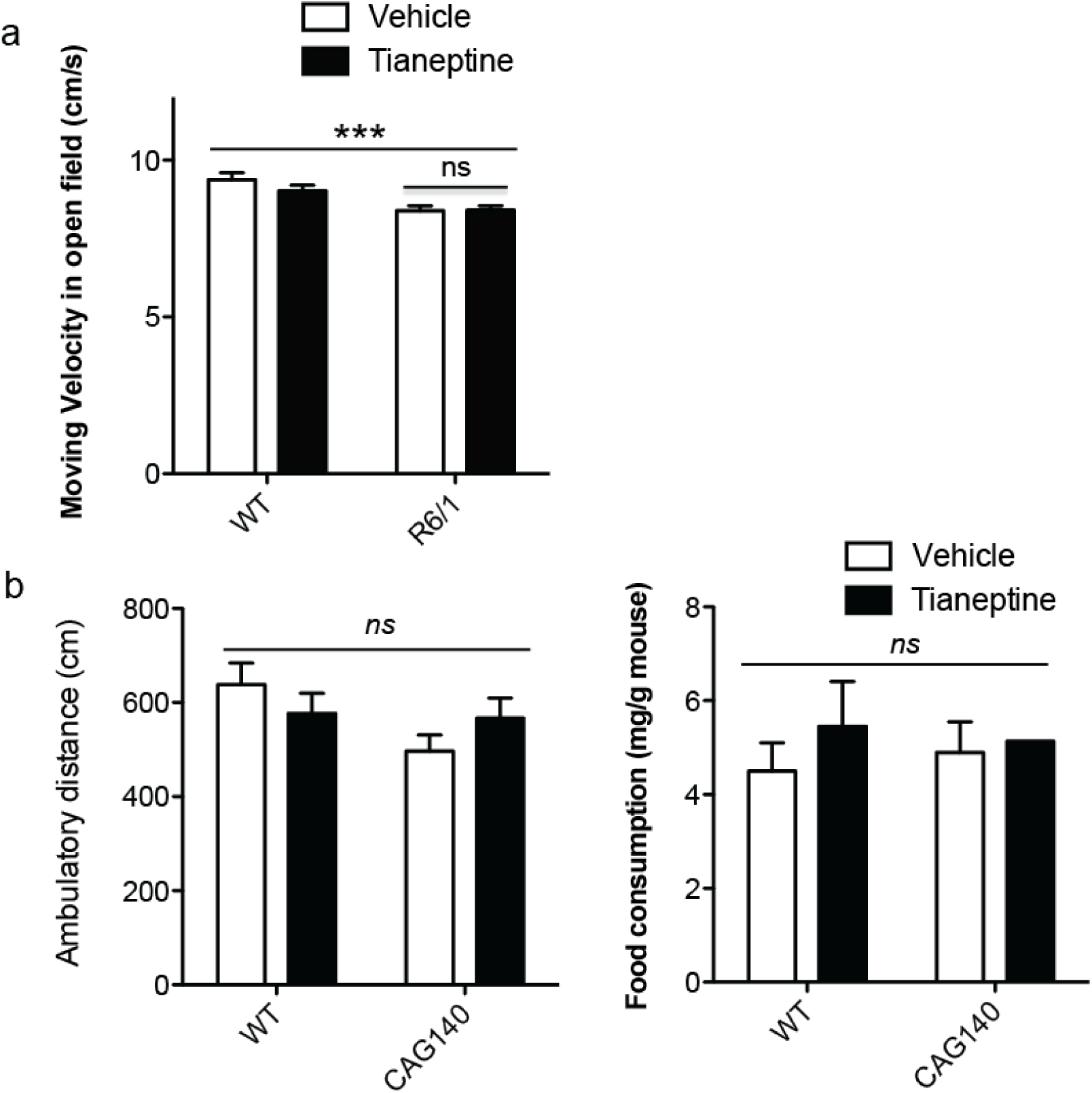
Tianeptine did not affect moving velocity of R6/1 mice in Open Field test, not change ambulatory distance nor food assumption of HTT CAG140 mice in elevated plus maze (EPM) and novelty-suppressed feeding (NSF), respectively. (**a**) Moving velocity in open field was significantly different between genotype but not between treatment; values are mean ± s.e.m; n = 25, 28, 33 and 32 mice for vehicle- and tianeptine-treated WT and R6/1 mice, respectively. (**b**) In EPM, there is no significant change in the locomotor activity between genotype nor treatment, which is revealed by ambulatory distance. (**c**) In NSF, food consumption was not significantly different between genotype nor treatment; values are mean ± s.e.m; n = 12, 9, 14, and 13 mice for vehicle- and tianeptine-treated WT and HTT CAG140 mice, respectively. Significance was assessed by two-way ANOVA followed by Bonferroni posttests (**a, b, c**). \*\*\**P* < 0.001; *ns*, not significant.

